# Comparative analysis of 16S rRNA gene and metagenome sequencing in pediatric gut microbiomes

**DOI:** 10.1101/2021.02.20.432118

**Authors:** Danielle Peterson, Kevin S. Bonham, Sophie Rowland, Cassandra W. Pattanayak, RESONANCE Consortium, Vanja Klepac-Ceraj

**Affiliations:** Department of Biological Sciences, Wellesley College, Wellesley, MA, USA; Department of Quantitative Reasoning and Mathematics, Wellesley College, Wellesley, MA, USA; Advanced Baby Imaging Lab, Hasbro Children’s Hospital, Rhode Island Hospital, Providence, RI, USA

**Keywords:** 16S rRNA gene, metagenome, pediatric cohort, gut microbiome, sequencing depth, amplicon sequencing

## Abstract

The colonization of the human gut microbiome begins at birth, and, over time, these microbial communities become increasingly complex. Most of what we currently know about the human microbiome, especially in early stages of development, was described using culture-independent sequencing methods that allow us to identify the taxonomic composition of microbial communities using genomic techniques, such as amplicon or shotgun metagenomic sequencing. Each method has distinct tradeoffs, but there has not been a direct comparison of the utility of these methods in stool samples from very young children, which have different features than those of adults. We compared the effects of profiling the human infant gut microbiome with 16S rRNA amplicon versus shotgun metagenomic sequencing techniques in 130 fecal samples; younger than 15, 15-30, and older than 30 months of age. We demonstrate that observed changes in alpha-diversity and beta-diversity with age occur to similar extents using both profiling methods. We also show that 16S rRNA profiling identified a larger number of genera and we find several genera that are missed or underrepresented by each profiling method. We present the link between alpha diversity and shotgun metagenomic sequencing depth for children of different ages. These findings provide a guide for selecting an appropriate method and sequencing depth for the three studied age groups.

## 1 Introduction

There is increasing evidence that changes in activity and diversity of the gut microorganisms are associated with the development of diseases and conditions such as type II diabetes (Hartstra et al., 2015; Lambeth et al., 2015), cancer (Bultman, 2014; Marchesi et al., 2011), and even depression (Foster and McVey Neufeld, 2013). Assessing the taxonomic diversity of gut microbes is a key first step towards understanding how those microbes may affect host health. Most of what is currently known about the gut microbiome has been derived using culture-independent profiling methods such as next-generation sequencing (Ji and Nielsen, 2015; Lozupone et al., 2012; Malla et al., 2019). The two most widely used culture-independent methods are amplicon sequencing, a method that amplifies variable regions of a highly conserved bacterial gene such as the 16S rRNA gene, and shotgun metagenomic sequencing, an approach that sequences all of the DNA present in a sample.

Both of these techniques have been pivotal in understanding the microorganisms living in the human gut and how they affect human health, but each has trade-offs. Profiling microbial communities using 16S rRNA genes is a straightforward and cost-effective method to profile the taxonomic composition of a microbial community, but it has low taxonomic resolution due to the conservation of the target gene and length of amplicon product. In addition, the amplification that is used to enrich for the rRNA gene can introduce bias in quantifying taxa in the resulting taxonomic profiles (Acinas et al., 2005; Tremblay et al., 2015). For instance, the choice of primers that bind to the 16S rRNA gene during amplification has been shown to have a great effect on microbiome community characterization (Chen et al., 2019; Tremblay et al., 2015). However, despite the need for a PCR amplification step, this type of profiling requires a relatively low number of sequenced reads per sample to maximize identification of rare taxa and is generally cheaper than shotgun metagenomic sequencing.

Shotgun metagenomics indiscriminately sequences the entire metagenome, and therefore typically requires more sequenced reads per sample to find unique taxonomic identifiers. This need for increased sequencing depth carries a higher cost (Comeau et al., 2017), but yields information on many genes rather than only one. This substantially increases resolution in taxonomic assignments - metagenomic profiling often provides species-level assignment where amplicon sequencing is restricted to identifying genera (Ranjan et al., 2016) - and has the additional benefit of providing direct evidence of gene functional variation in strains present. Metagenomic sequencing may also be used to generate genomic assemblies, yielding further insight into microbial diversity (Wilkins et al., 2019).

The ability to draw conclusions about taxonomy from microbiome sequencing data depends not only on the sequencing method, but also on sequencing depth: how many times on average a given piece of DNA is likely to be sequenced given a fixed read length and the assumption that all regions of a genome are equally likely to be sequenced (Sims et al., 2014). If it were possible to achieve the resolution of shotgun metagenomics at a lower cost, we could sequence more deeply, identify less abundant taxa, and learn more about the microbial diversity within and between samples (Pereira-Marques et al., 2019). However, deeper sequencing is more expensive. A few studies have investigated the potential for reduced metagenomic sequencing (Hillmann et al., 2018; Zaheer et al., 2018), but there has not been substantial research analyzing the reduced sequencing depth for investigation of the gut microbiomes in young infants and children. The gut microbial communities of children are potentially good candidates for experimentation with shallower sequencing depths because their communities have lower gut microbial diversity until their microbiomes stabilize and become more adult-like around 2-3 years of age (Palmer et al., 2007; Radjabzadeh et al., 2020; Stewart et al., 2018; Yatsunenko et al., 2012).

While some have utilized both profiling methods in children (Ravi et al., 2018; Vatanen et al., 2016), and the known trade-offs between amplicon and metagenomic sequencing have been previously explored in soil (Brumfield et al., 2020) and plant environments (Mas-Lloret et al., 2020), as well as in human adult microbiomes (Laudadio et al., 2018; Ranjan et al., 2016), to date, no one has directly investigated the relative trade-offs between 16S rRNA amplicon sequencing and metagenomic sequencing at different sequencing depths in the gut microbiomes of infants and young children of different ages. Here, we compare paired 16S rRNA versus metagenomic sequencing gut microbiome datasets from a cohort of young children broken into 3 age brackets: less than 15, 15 to 30, and over 30 months.

## 2 Materials and Methods

### 2.1 Cohort description

Samples for this study came from a subset of 130 children (**Figure S1**) in the RESONANCE Cohort (Providence, RI), an accelerated-longitudinal study of healthy children ages 0 to 12 years. Each child contributed one sample. The RESONANCE cohort is part of the <u>E</u>nvironmental influences on <u>C</u>hild <u>H</u>ealth <u>O</u>utcomes (ECHO) Program (Forrest et al., 2018; Gillman and Blaisdell, 2018), which aims to investigate the effects of environmental factors on childhood health and development. All procedures for this study were approved by the local institutional review board at Women and Infants Hospital, and all experiments adhered to the regulation of the review board. Written informed consent was obtained from all parents or legal guardians of enrolled participants. Children with known major risk factors for developmental abnormalities at enrollment were excluded.

### 2.2 Stool sample collection and handling

One stool sample per child (n=130) was collected by parents in OMR-200 tubes (OMNIgene GUT, DNA Genotek, Ottawa, Ontario, Canada), stored on ice, and brought within 24 hrs to the lab in RI where they were immediately frozen at −80°C. Stool samples were not collected if the infant had taken antibiotics within the last two weeks. Samples were transported to Wellesley College (Wellesley, MA) on dry ice for further processing.

### 2.3 DNA extraction and sequencing of metagenomes and 16S rRNA gene amplicons

Nucleic acids were extracted from a 200 μL aliquot of fecal slurry using the RNeasy PowerMicrobiome kit automated on the QIAcube (Qiagen, Germantown, MD), according to the manufacturer’s protocol, excluding the DNA degradation steps. The samples were subjected to bead beating using the Qiagen PowerLyzer 24 Homogenizer (Qiagen, Germantown, MD) at 2500 speed for 45 seconds. The samples were transferred to the QIAcube to complete the protocol, and extracted DNA was eluted in a final volume of 100 μL. DNA extracts were stored at −80°C until sequenced.

Samples were sequenced at the Integrated Microbiome Resource (IMR, Dalhousie University, NS, Canada) (Comeau et al., 2017). To sequence metagenomes, a pooled library (max 96 samples per run) was prepared using the Illumina Nextera Flex Kit for MiSeq and NextSeq (a PCR-based library preparation procedure) from 1 ng of each sample where samples were enzymatically sheared and tagged with adaptors, PCR amplified while adding barcodes, purified using columns or beads, and normalized using Illumina beads or manually. Samples were then pooled onto a plate and sequenced on the Illumina NextSeq 550 platform using 150+150 bp paired-end “high output” chemistry, generating ~400 million raw reads and ~120 Gb of sequence (NCBI Bioproject PRJNA695570).

For sequencing 16S rRNA gene amplicons, the V4-V5 region of the 16S ribosomal RNA gene was sequenced according to the protocol described by Comeau et al. (2017). Briefly, the V4-V5 region was amplified once using the Phusion High-Fidelity DNA polymerase (ThermoFisher Scientific, Waltham, MA) and universal bacterial primers 515FB: 5’-GTGYCAGCMGCCGCGGTAA-3’ and 926R: 5’-CCGYCAATTYMTTTRAGTTT-3’ (Parada et al., 2016; Walters et al., 2016). These primers had appropriate Illumina adapters and error-correcting barcodes unique to each sample to allow up to 380 samples to be simultaneously run per single flow cell. After being pooled into a single library and quantified fluorometrically, samples were cleaned-up and normalized using the high-throughput Charm Biotech Just-a-Plate 96-well Normalization Kit (Charm Biotech, Cape Girardeau, MO). The normalized samples were sequenced on the Illumina MiSeq platform (Illumina, San Diego, CA) using 300+300 bp paired-end V3 chemistry, producing ~55,000 raw reads per sample (Comeau et al., 2017).

### 2.4 16S rRNA gene amplicon processing and analysis

Reads profiled using the 16S rRNA gene were analyzed using the Quantitative Insights in Microbial Ecology 2 (QIIME2), v 2019.10 (Bolyen et al., 2019) and we used a modified protocol developed by Comeau et al. (2017). Briefly, primers flanking V4-V5 were removed from fastq reads using the cutadapt QIIME2 plugin (Martin, 2011). Fastq reads were then filtered, trimmed and merged in DADA2 (Callahan et al., 2016) to generate a table of amplicon sequence variants (ASV). A multiple-sequence alignment was created using MAFFT, and FastTree was used to create an unrooted phylogenetic tree, both with default values (Price et al., 2010). A root was added to the tree at the midpoint of the largest tip-to-tip distance in the tree. Taxonomy was assigned to the ASVs using a Naïve-Bayes classifier compared against a SILVA v 119 reference database trained on the 515-926 region of the 16S rRNA gene (Bokulich et al., 2018). Rarefaction curves showed that the majority of samples reached asymptote, indicating sequencing depth was appropriate for analyses.

### 2.5 Metagenome data processing and analysis

Metagenomic data were analyzed using bioBakery workflows with all necessary dependencies and default parameters (McIver et al., 2018). Briefly, KneadData (v 0.7.10) was used to trim and filter raw sequence reads, and to separate human and 16S ribosomal gene reads from bacterial sequences in both fecal and oral samples. Samples that passed quality control were taxonomically profiled to the genus level using MetaPhlAn (v 3.0.7), which uses alignment to a reference database of “marker genes” to identify taxonomic composition (Beghini et al., 2020).

### 2.6 Statistical Analysis

Statistical analyses were carried out in R (4.0.3). *vegan* (v 2.5-6) was used for all alpha-diversity calculations: Shannon diversity index (Shannon, 1948) (alpha diversity measurement of evenness and richness), evenness (how homogeneous the distribution of taxa counts are), and richness (number of taxa in a community). Pairwise Bray-Curtis dissimilarity was used to assess beta-diversity, or the overall variation between each sample (Bray and Curtis, 1957). The Bray-Curtis dissimilarity metric compares two communities based on the number or relative abundance of each taxon present in at least one of the communities. When we calculated these values, we assumed that the set of dissimilarities calculated across a group was independent, even when the same child was paired to other children multiple times. These distance matrices were used for Principal Coordinates Analysis (PCoA) to create ordinations. The two principal components that explained the most variation were used to create biplots (**Figure S2**).

Univariate comparisons were performed in two-sample two-tailed t-tests when we could assume normality, and Wilcoxon Signed Rank tests when we could not. P-values of less than 0.05 (or the equivalent after Benjamini-Hochberg false discovery rate correction (Benjamini and Hochberg, 1995)) were considered statistically significant. Mixed effects linear models in *lme4* were used to analyze data from subsampling results, to account for the fact that multiple subsamples were generated from each sample. Shannon ~ 1.58 + 5.21×10^−4^*read depth - 3.79×10^−1^ *less than 15 months - 4.38×10^−2^ *older than 30 months - 1.50×10^−4^*read depth: less than 15 months - 1.56×10^−4^*read depth:older than 30 months.

### 2.7 Comparing missing and underrepresented genera in 16S rRNA to shotgun metagenomics datasets

A genus was classified as being unique to a particular profiling method if reads were only assigned to it through one method. Taxa that could not be resolved down to the genus level (taxonomic assignments containing the phrases “unclassified,” “unidentified,” “group,” or “uncultured”) were removed prior to calculating relative abundance diversity, and all downstream metrics. Genera that only occurred in one but not the other method were classified as uniquely identified by 16S rRNA profiling or shotgun metagenomics. We found the intersection of genera by identifying microbes that were found at least once by both methods.

We used Wilcoxon Signed Rank tests to compare the abundances of microbes that were found by both methods. This analysis was limited by the direct comparison of relative abundances instead of direct counts. Because 16S rRNA profiling was able to identify more taxa at the genus level, this meant that the relative abundances of its organisms were systematically lower.

### 2.8 Analyzing primer coverage

TestPrime 1.0 (Klindworth et al., 2013; Ludwig et al., 2004) was used to perform *in silico* PCR to investigate how well certain primer pairs align to microbes in the SILVA database. We entered our forward and reverse primers (515FB and 926R) into the TestPrime web-tool provided by SILVA (Quast et al., 2013) to analyze the percent primer coverage of microbes found only with metagenomic sequencing, but not by amplicon sequencing. Coverage is defined as the percentage of matches for a particular taxonomic group (# of matches / (total # of mismatches + matches). The primers described in Methods 2.4 were compared to sequences found within the SSU r138.1 SILVA database. A single nucleotide mismatch between each primer and 16S rRNA gene sequence was considered a mismatch for that organism. Once the percent coverage was calculated, we compared the average coverage of microbes uniquely found by shotgun metagenomics, 16S rRNA profiling, or both methods. Some genera identified uniquely by shotgun metagenomics were not as identified as hits to the primer, despite being in the SILVA database. Their alignment was manually entered to be 0% for downstream analysis.

### 2.9 Generating phylogenetic trees

The union of all genera that were identified by either 16S rRNA gene or shotgun metagenomic sequencing was used to generate a phylogenetic common tree using TimeTree (Kumar et al., 2017). In addition to these genera, *Thermus aquaticus* was added as an outgroup. This tree was visualized using the Interactive Tree of Life (iTOL) v 5.5.1, (Letunic and Bork, 2019), along with metadata that described which profiling method (either 16S, shotgun metagenomics, or both) was able to identify the genus (Letunic and Bork, 2007, 2019). For taxa that were unidentified by a particular profiling method, we investigated whether or not that taxon was present in the missing database. The phylogenetic tree notes taxa that would be impossible to be identified by that method, as they were not present in the relevant database.

### 2.10 Exploring the effect of read depth on diversity using metagenome samples

We investigated the results of decreasing read depth on alpha and beta-diversity by resampling shotgun metagenomic reads from a subset of children within the RESONANCE cohort that had deeply sequenced metagenomes (average 7,209,871± 2,562,647 reads). Metagenomic reads from 30 children were selected and 10k, 100k, 250k, 500k, 750k, and 1M reads were randomly sampled (with replacement) from each child’s reads. Each child was resampled at each depth four times for the analysis involving RESONANCE subjects.

To investigate whether these observations were generally applicable to other childhood cohorts, we performed the same subsampling analysis on the DIABIMMUNE cohort (Simre et al., 2016). Only a single sample for each depth was obtained for DIABIMMUNE subjects due to the substantially higher number of original samples. DIABIMMUNE subjects were subsampled at depths of 100k, 250k, 500k, 750k, 1M, and 10 M reads. All children were separated by developmental stage (less than 15 months: n = 10, between 15 and 30 months, n = 10, over 30 months, n = 10). Reads were reassigned taxonomy using MetaPhlAn (see section 2.7) and diversity was recalculated. The majority of these samples subsampled at 10,000 reads had no identifiable taxa and were excluded from downstream analysis.

## 3 Results

### 3.1 Alpha diversity increases with age in both 16S rRNA gene- and metagenomic-profiled samples

First, we directly compared taxonomic profiles generated by shotgun metagenomic or amplicon sequencing to assess their ability to detect poorly characterized or low abundance taxa. On average, the proportion of microbes resolved to the genus level in a sample was 97.7% (SD = 1.7%) when profiled by shotgun metagenomic sequencing and 78.2% (SD = 20.7%) when profiled by 16S rRNA sequencing. As expected, regardless of the profiling method, the observed alpha (within-sample) diversity of the gut microbiome of children increased in the first 30 months of life (Welch’s t-test, p-value < 0.001). Given that we observed that children’s microbiomes grow increasingly complex and diverse, we hypothesized that any differences in ability of the profiling methods to identify less-abundant taxa would only be magnified with age. Consistent with this hypothesis, we found that profiles created from shotgun metagenomics data had systematically lower alpha diversity than profiles from 16S rRNA sequencing at the genus level across all developmental stages (**Figure 1A**). The mean of these differences between paired profiles increased as the children age, with the largest differences observed in children older than 30 months (mean of the differences = 0.18, paired t-test, p-value < 0.001). This suggests that the differences between 16S rRNA and shotgun metagenomics profiling in capturing alpha diversity are amplified as children age and their microbial diversity becomes increasingly complex.

**Figure 1:**
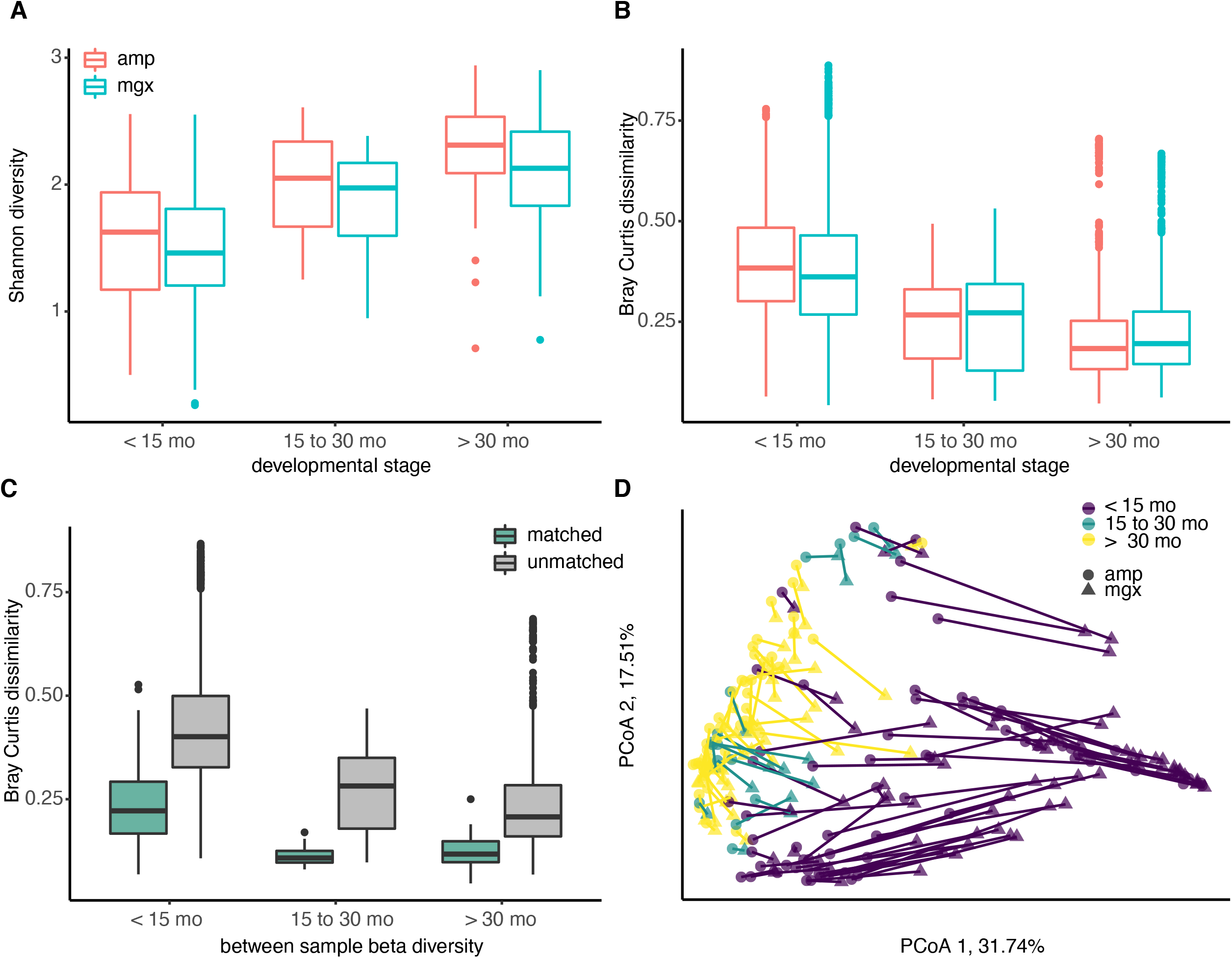
Diversity of the child gut microbiome differs by age, regardless of profiling method. Microbiome communities from 130 children were sequenced using 16S rRNA (abbreviated “amp”) and shotgun metagenomic (abbreviated “mgx”) profiling. **(A)** Alpha diversity was calculated using the Shannon diversity index for each child. Boxplots are grouped by age and colored by profiling method. **(B)** Beta-diversity was quantified using pair-wise Bray-Curtis dissimilarities between all children within the same profiling method and developmental stage. **(C)** Bray-Curtis dissimilarities between 16S and metagenomic profiles for matched samples (from same fecal sample), 16S and metagenomic profiles among unmatched samples (from different fecal samples). **(D)** Beta-diversity was visualized using Principal Coordinate analysis (PCoA). The first two principal coordinate axes, which together explain 49.25% of variation, are shown. Each dot represents one taxonomic profile, with lines connecting profiles from the same sample. Colors represent developmental stages and shape represent profiling methods.

We next examined between-sample, or beta, diversity within each of the three age groups to determine if age or profiling method were associated with large between-sample differences. Comparisons of beta diversity within children of the three groups indicated the similarity between gut microbiome communities increased with age in both profiling methods. Regardless of which method was used, Bray-Curtis dissimilarity, a pairwise measure of beta diversity between two communities, was the smallest between children over the age of 30 months (**Figure 1B**).

After observing differences in the two profiling methods among young children, we next compared profiles generated from the different methods for the same fecal sample. If data from shotgun metagenomics and 16S rRNA gene profiling both produced exactly the same gut microbial profiles, we would expect that profiles from the same child’s fecal sample would have a Bray-Curtis dissimilarity of ~0. At a minimum, we would expect to see that the Bray-Curtis dissimilarity among profiles constructed from the same stool sample would be smaller than the dissimilarity between two profiles from two random children. As hypothesized, we observe that the average Bray-Curtis dissimilarity among paired samples is much lower than that of unpaired samples (**Figure 1C;** mean difference = 0.348, Welch’s t-test, p-value < 0.001). The largest differences in the paired profiles were found in children less than 15 months (**Figure 1C, 1D**).

### 3.2 Discrepancies between 16S rRNA and shotgun metagenomics profiles

To further investigate the cause of the largest discrepancies in diversity between the two profiling methods, we looked at biases in taxonomic representation at different taxonomic levels. At all taxonomic levels, except the species level, 16S rRNA amplicon profiling identifies more taxa (Figure 2A). We found that 41 families were found by both methods, while 33 and 14 were uniquely identified by 16S rRNA and shotgun metagenomic profiling, respectively. At the genus level, of 202 genera identified across all samples, only 105 genera were identified with both amplicon and shotgun metagenomic sequencing. 16S rRNA amplicon sequencing identified 63 genera not found by metagenomic profiling including *Acetobacter*, *Bacillus*, *Flavobacterium, Pseudomonas*, and *Sulfitobacter*, while only 34 genera were uniquely found using shotgun metagenomic sequencing, such as *Citrobacter*, *Coprococcus*, *Enterobacter, Gordonibacter,* and *Helicobacter* (Figure 2B). At the species level, 16S rRNA amplicon profiling was not able to resolve any taxa to the species level, while shotgun metagenomics was able to identify 385 unique species. We decided to focus on comparing taxonomic differences at the genus level, as that is the most specific taxonomic level in which we are able to meaningfully compare the two methods.

**Figure 2:**
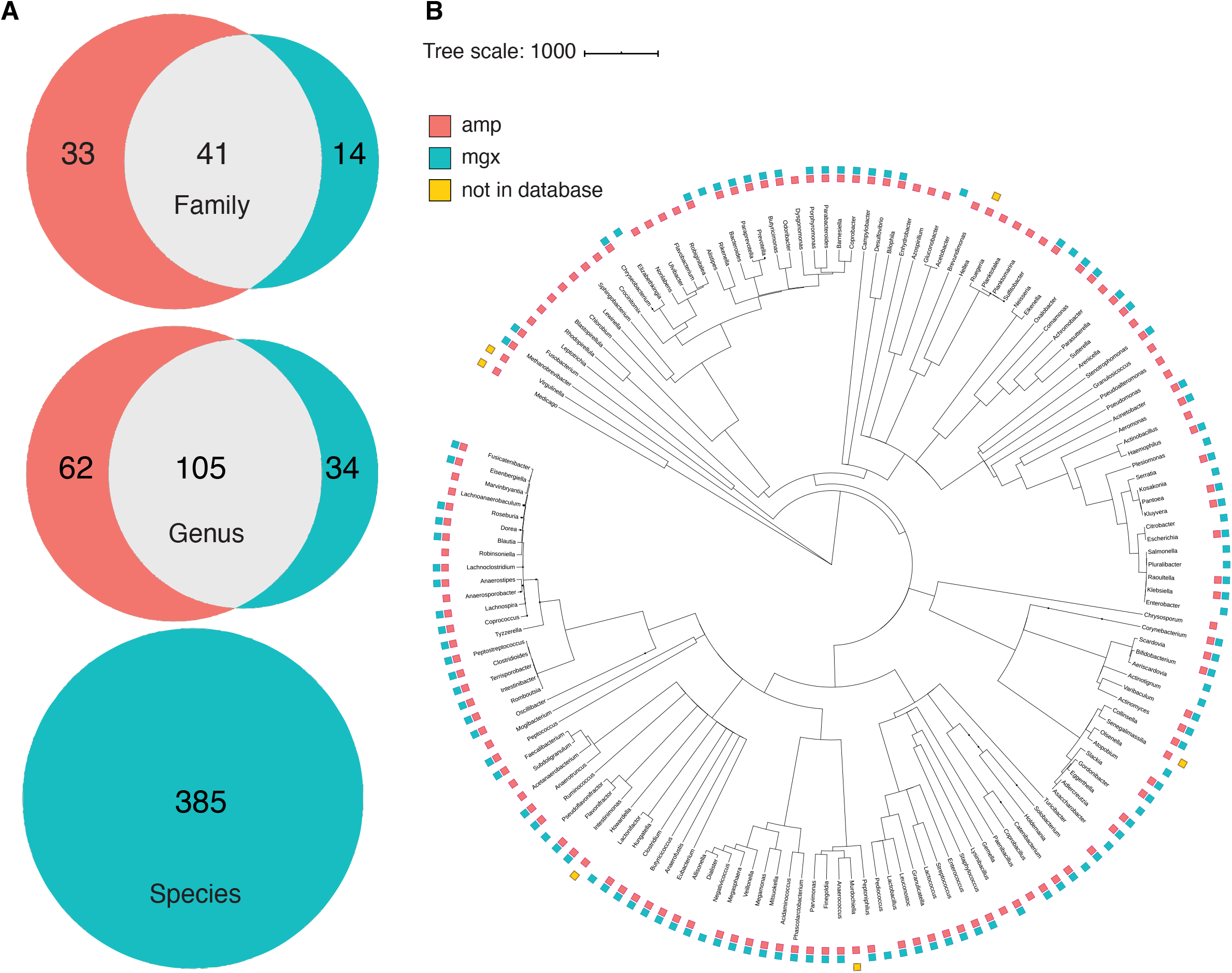
Some phylogenetic clustering of taxa by profiling method. **(A)** Venn diagrams indicating the number of taxa that were found by 16S (peach), shotgun metagenomics (cyan), or both (grey) methods. Number of overlapping and unique taxa were calculated on the family, genus, and species level. **(B)** A common phylogenetic tree was generated from all taxa identified by both 16S rRNA gene (amp) and shotgun metagenomic sequencing (mgx). Colors indicate which method was able to identify taxa (peach = identified by 16S, cyan = identified by shotgun metagenomics, yellow = taxa was not present in the database of the method with which it was not found).

After identifying genera that were found by only one of the two methods, we next investigated whether there were any taxa that were systematically found at higher levels in one method versus the other. We found that *Butyricicoccus* was observed to have a significantly higher relative abundance in 16S rRNA profiles compared to samples profiled with shotgun metagenomics (Wilcoxon signed rank test for this and all microbes, p-value < 0.001) (**Table S1**). Similarly, *Romboutsia* (p-value < 0.001) and *Sutterella* (p-value < 0.001) were found to have a higher relative abundance when detected by 16S rRNA amplicon sequencing. In contrast, genera such as *Bifidobacterium* (p-value < 0.001)*, Eggerthella* (p-value < 0.001), and *Klebsiella* (p-value < 0.001) systematically had higher relative abundance when detected by shotgun metagenomic techniques.

### 3.3 Reduced sequencing depth decreases has smaller effect on observed diversity in young children

After comparing two different profiling methods, we investigated the effect of reducing metagenomic sequencing depth on observed alpha diversity among the three developmental groups. We selected samples from a sub-group of 30 children (10 from each developmental stage) that were initially sequenced at the highest depth (mean 7.2 million reads; SD = 2.6 million reads) and performed random resampling of shotgun metagenomic reads at varying depths (100k, 250k, 500k, 750k and 1M reads). We then recalculated alpha diversity metrics (evenness, richness, and Shannon) for each community of re-sampled reads after assigning taxonomy using MetaPhlAn. **Figure 3A** shows the relationship between the evenness, richness, and sequencing depth across all the resamplings we performed. Regardless of the starting community’s diversity, as sequencing depth increased, observed sample richness and evenness also increased (**Figure S3**). For example, samples that were only profiled with 100k reads had a mean Shannon Index of 1.35, whereas those sampled at 1M reads had mean Shannon Index of 1.89 (**Figure 3B**).

**Figure 3.**
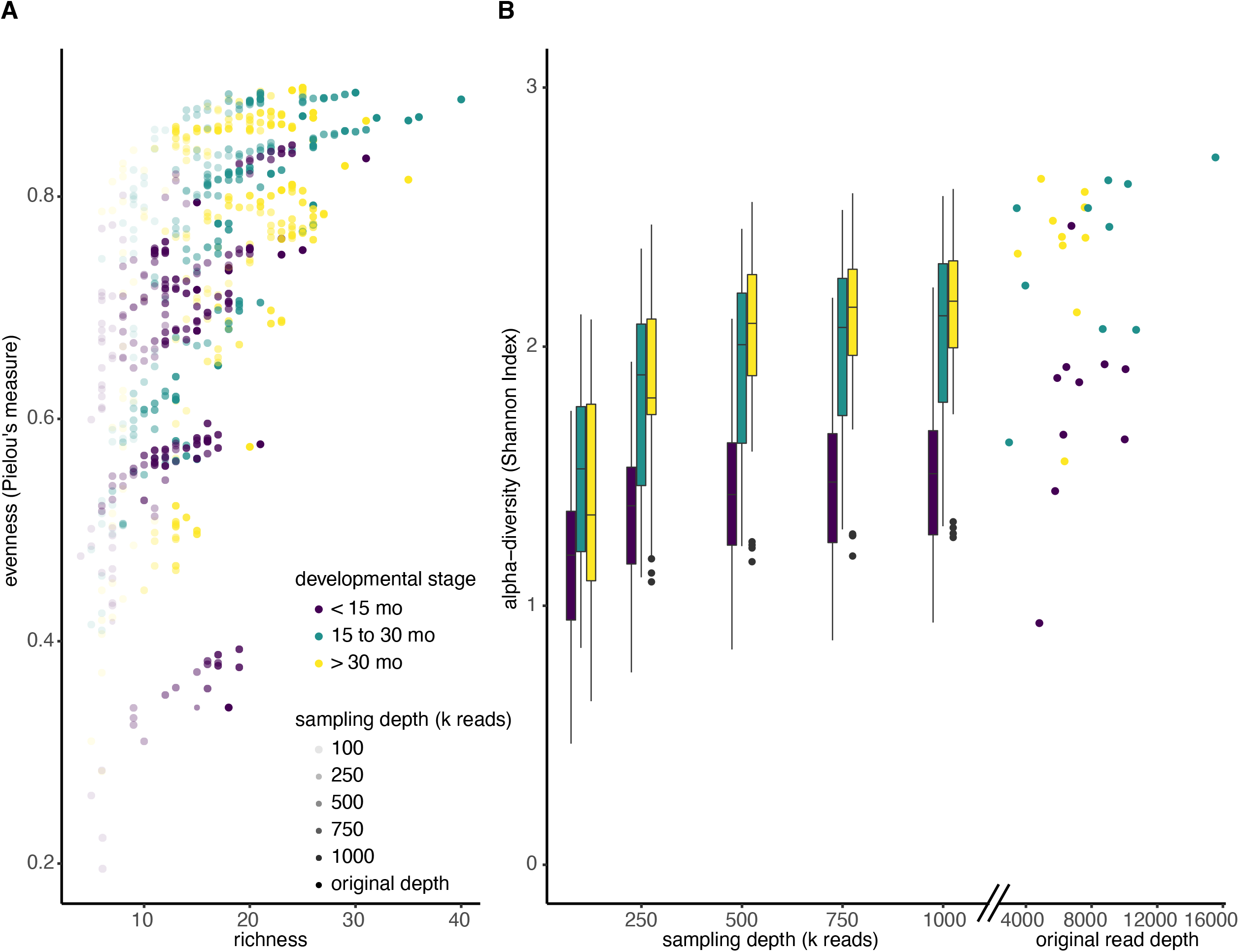
Alpha diversity increases with sequencing depth. **(A)** Shotgun metagenomic reads from 30 deeply sequenced samples were resampled four times at each different sequencing depths (100k; 250k; 500k; 750k; 1M reads). Reads were reassigned taxonomy using MetaPhlAn and diversity was recalculated. Each dot represents a single resampled community. **(B)** Boxplots of Shannon diversity among all samples at each re-sampling depth, colored by developmental stage. Scatter plot indicates Shannon diversity of original samples.

In addition, we observed that increasing sequencing depth affected children of different ages differently. Not only did children younger than 15 months have a lower median Shannon Index when we ignore sampling depth (<15 months median: 1.42, >15 months median: 1.99), the Shannon Index increases more slowly with sampling depth in kids under 15 months. In particular, a mixed effects linear model showed that the slope of the Shannon Index on sampling depth is significantly lower for children under 15 months, compared to those between 15 and 30 months (p < 0.001), and the slope is significantly lower for children between 15 and 30 months compared to those greater than 30 months (**Figure 3B**; p < 0.001).

While the step-wise increase in alpha diversity with sampling depth is statistically significant for children less than 15 months (p < 0.001), the increase in observed alpha diversity is substantially smaller than typical effect sizes in childhood microbiome studies. For instance, a recent meta-analyses of other studies that investigated alpha diversity of children that were and were not breastfed observed average differences in Shannon Index to be 0.34 (95% Confidence Interval: [0.20, 0.48]) (Ho et al., 2018), but increasing sequencing depth from 500k reads to 1M reads only increased this metric by 0.06 (**Table 1**, **Table S2**).

**Table 1.** Average Shannon Index values among children less than 15 months at different subsampling depths in the RESONANCE data-set. “Read_depth”: indicates subsampling depth, “mean”: mean Shannon Index at subsampling depth, sd: standard deviation of Shannon Index at subsampling depth, “abs_diff”: absolute difference in Shannon Index from previous subsampling depth.

| **read_depth** | **mean** | **sd** | **abs_diff** |
| --- | --- | --- | --- |
| **100** | 1.35 | 0.39 |  |
| **250** | 1.67 | 0.42 | 0.32 |
| **500** | 1.8 | 0.44 | 0.13 |
| **750** | 1.86 | 0.44 | 0.06 |
| **1000** | 1.89 | 0.44 | 0.03 |
| **original** | 2.04 | 0.42 | 0.15 |

To investigate whether these observations were generally applicable to other childhood cohorts, we performed the same subsampling analysis on the DIABIMMUNE cohort (Simre et al., 2016) (**Figure S4**). Consistent with the findings from the RESONANCE cohort, lower sequencing depth decreases the Shannon Index for all age groups (Mixed effects linear model, p <0.001), and the benefits of deeper sequencing are most pronounced in older kids, as observed alpha diversity increases more quickly as additional reads are added for older children (**Table S3**, p < 0.001). In addition, for both cohorts, the benefits of additional sequencing on observed diversity in children under 15 months substantially decrease over 500 thousand reads.

## 4 Discussion

Increasing interest in the human microbiome, especially during early child development, raises the urgency of selecting appropriate methods for interrogating taxonomic and functional composition of human-associated communities. Given that shotgun metagenomic sequencing is capable of providing higher taxonomic resolution as well as information about gene functional potential, it is clearly preferable to amplicon sequencing when working with high biomass samples such as stool and when cost is not an issue. However, the higher cost of sequencing to provide sufficient sequencing depth for shotgun metagenomics is relevant when resources are constrained. Because infant microbiomes are substantially less diverse than adult microbiomes, we reasoned that lower sequencing depth (and therefore lower cost) may enable comparable taxonomic resolution to amplicon sequencing at a similar cost.

We, therefore, set out to analyze a group of child stool samples sequenced with both methods and profiled with commonly used taxonomic-assignment tools so that direct comparisons could be made. As expected, microbial communities from younger children (less than 15 months old) were substantially less diverse than communities from older children, and both amplicon and shotgun metagenomic sequencing with ~1.2 Gb per sample were able to capture comparable taxonomic diversity at the genus level across all age groups. It is important to note that metagenomic sequencing generally captures more diversity due to its species-level resolution (Ranjan et al., 2016), but we restricted our analysis to the genus level in order to make the most direct comparison to amplicon sequencing. Interestingly, though the observed diversity overall was comparable between methods, the actual taxonomic profiles generated by each method had substantial differences, particularly in the youngest children. For example, some particularly important genera in young children such as *Bifidobacterium* and *Enterobacter* were under-represented in amplicon sequencing profiles. Because shotgun metagenomic sequencing does not include an amplification step and therefore avoids issues of amplification bias, it is likely to be more accurate, though further investigation with synthetic or *in silico* communities may be necessary to determine which method provides the most accurate profiles in this population.

While shallower sequencing may enable investigators to observe comparable diversity, there are substantial differences in the identities of taxa profiled. Like other groups (Rausch et al., 2019), we showed that 16S rRNA gene amplicon and shotgun metagenomic sequencing each missed some taxa, but more genera were identified overall by 16S rRNA gene profiling, at least in the RESONANCE cohort. This may be due to an increased ability to identify very low abundance taxa or some artifact of amplification or sequencing, though in the DIABIMMUNE cohort, more genera were identified using shotgun metagenomic profiling, suggesting that the relative performance of each method for some metrics may vary between populations. Interestingly, we also show that the largest discrepancies between the two profiling methods were found in the youngest kids. This is likely due in part to the low diversity of these samples, since loss of one genus in a profile with few genera may have a larger impact on dissimilarity metrics. Another possible explanation is the large fraction of many samples in young children (as much as 40% relative abundance) that could not be resolved to the genus level (see section 3.1) with amplicon sequencing. As unresolved taxa were excluded from our alpha diversity analysis, the true diversity could be much higher or lower than we observe in those samples.

Some of the discrepancies we observed were due to technical differences in sequencing methods. For example, some taxa found exclusively through 16S rRNA gene profiling were not found in the MetaPhlAn database, including 16 genera that did not have reference genomes available. All of the genera found uniquely by shotgun metagenomics were present in the SILVA database, but their 16S rRNA gene sequences may not have perfectly complemented the primers we used. Though 16S rRNA PCR primers are often referred to as “universal,” there is considerable sequence diversity in the 16S rRNA gene, even in the most conserved regions and among bacteria of the same species (Větrovský and Baldrian, 2013). Using TestPrime 1.0, we identified several genera that had very low alignment with our primers, such as *Solobacterium* (2.2% alignment) and *Pediococcus* (1.3%) and 10 genera that were present in the SILVA database and identified using shotgun metagenomics, but were not found to be hits with our primers. We also explored if certain clusters of taxa were more systematically unidentified by a particular profiling method. For example, several genera identified uniquely by shotgun metagenomic profiling had lower primer coverage compared to the genera identified by 16S rRNA amplicon profiling (**Figure S5**). Other taxa were only identified using 16S amplicon profiling (**Figure 2B**; ex. clade containing *Ruegeria*, *Planktotalea*, *Planktomarina*, and *Sulfitobacter*).

Given that both profiling methods exhibited some biases against certain taxa, future study designs should carefully consider which method is most appropriate to their research question, and further investigation using communities where the ground truth of composition is known should be pursued to interrogate whether these differences are systematic. In addition to uncertainty about the true composition of these samples’ communities, our study was also limited in scope to a single 16S rRNA gene primer pair for amplification, a single sequencing read length for shotgun sequencing, and a single computational pipeline for taxonomic profiling each sequencing method. There are several different approaches for both the sequencing (Driscoll et al., 2017; Martínez et al., 2014; Rausch et al., 2019) and profiling step (Almeida et al., 2018; Ye et al., 2019), each of which is likely to have its own biases. We chose to compare widely used and accessible methods to compare for investigation of child microbiomes, but further investigation to select the best combination of methods may be warranted. Finally, advances in sequencing technology (e.g., long-read sequencing of 16S rRNA genes (Karst et al., 2020)), changes to reference databases and improved taxonomic assignment methods may affect the performance and relative trade-offs in the future.

## 5 Conclusion

Understanding the advantages associated with different methods of investigating the human microbiome will allow others in the field to use the most cost-effective methods to explore the relationship between the gut microbiome and human health. Most research is limited by financial resources, which impacts the number of controls, replicates, samples we can analyze, and the depth to which we can characterize each sample. Better insight into how we can sequence more efficiently will allow us to use these finite resources more effectively. Hopefully, this will allow us to devote resources where they will be best utilized (eg. deep sequencing for older children with higher alpha diversity) and reduce them where they are not necessary.

Given the importance of the first thirty months of one’s life in shaping future health outcomes (Bokulich et al., 2016; Tamburini et al., 2016; Yang et al., 2016), it is crucial that we understand how to efficiently characterize developing microbiomes. By identifying the most effective methods for investigating the microbiomes of children at different stages of development, we can reduce sequencing costs and reduce bias in results. This will ultimately increase the quality of the research by ensuring that resources are appropriately expended. Altogether, understanding the links between the infant gut microbiome and child development will allow us to better predict how early-life environmental exposures or health decisions can mediate the gut microbiome’s effects on health later in life.

## Author Contributions

DP, VKC, and KSB designed the study; SR processed the samples; DP and KSB designed and wrote the code and analyzed the data; CP contributed to data analyses and statistical methods. All authors wrote, edited, and finalized the manuscript. All authors approved the manuscript’s final version.

## Acknowledgments

We thank families participating in the RESONANCE cohort. We thank Maureen Coleman and Steven Biller for reviewing the manuscript and providing helpful comments, and Christopher Loiselle and Jennifer Beauchemin for assistance with biospecimen and metadata collection. The project was funded by the NIH UG3 OD023313 (VKC).

## Data and Code availability

Raw and processed data is available through SRA (NCBI Bioproject PRJNA695570) and at OSF.io (Peterson et al., 2021).

## Supplementary Materials

The supplementary material for this manuscript can be found online at:

## Supplemental Figures

**Figure S1: RESONANCE: a cohort of healthy children between ages 2 months and 4 years**

Histogram showing distribution of ages across developmental stages. Both 16S rRNA gene data and metagenome profiles were obtained for 130 stool samples (one sample per child and timepoint). n= 85 for children <15 months, n=15 for children 15-30 months, n= 60 for children >30 months. Color indicates developmental stage.

**Figure S2: Cumulative percent of variation explained by first 100 principal components**

Barplot of the cumulative sum of the percentage explained by the first 100 principal components used to create Figure 1D. The first 10 principal components explained 88.5% of the total variation in Bray-Curtis dissimilarity within the dataset.

**Figure S3: Species richness increases with sampling depth within developmental stage**

**(A)** Boxplots of species richness among all samples at each sampling depth, colored and grouped by developmental stage. (**B)** Boxplots of species richness among all samples at each re-sampling depth, separated by sampling depth.

**Figure S4: Alpha diversity decreases with sequencing depth in DIABIMMUNE dataset**

**(A)** Shotgun metagenomic reads from 804 deeply sequenced samples were resampled times at six different sequencing depths (100k; 250k; 500k; 750k; 1M, & 10 M reads). Reads were reassigned taxonomy using MetaPhlAn and diversity was recalculated. Each dot represents a single resampled community. **(B)** Boxplots of Shannon diversity among all samples at each re-sampling depth, colored by developmental stage. Scatter plot indicates Shannon diversity of original samples.

**Figure S5: Genera found by 16S rRNA amplicon sequencing have significantly higher primer coverage**

TestPrime 1.0 was used to calculate the percent primer coverage of the primers used in our study for amplicon sequencing. We compared the percent coverage for microbes found uniquely by 16S rRNA sequencing, both methods, and shotgun metagenomic sequencing. A pairwise Wilcoxon test found that primer coverage for microbes found uniquely by amplicon sequencing is significantly higher than that in the genera found uniquely by shotgun metagenomics (p < 0.05).

## Supplemental Tables

**Table S1: Genera systematically over-represented with either profiling method**

The Wilcoxon signed-rank test was used to compare the relative abundances of a particular genera, calculated from 16S and shotgun metagenomics profiling. “Diff” is the average relative abundance difference for a particular genera (mean 16S relative abundance - mean shotgun metagenomics relative abundance) “P.adjust” is the p-value after Benjamini-Hotchberg correction. The table presents genera with significant differences (adjusted p-value < 0.05), indicating genera that had higher average relative abundances when profiled by 16S rRNA or shotgun metagenomics. “Method” indicates the profiling method where the genus was more abundant.

**Table S2: Output of linear model used to predict Shannon Index based on read depth and developmental stage in RESONANCE dataset**

*lme4* was used to construct a Mixed effects linear model to analyze data from the RESONANCE subsampling results. “Estimates” reports the estimated coefficients for the intercept of the fitted line and each variable (read depth, developmental stage) or interaction of variables.

**Table S3: Output of linear model used to predict Shannon Index based on read depth and developmental stage in DIABIMMUNE dataset**

*lme4* was used to construct a Mixed effects linear model to analyze data from the DIABIMMUNE subsampling results. “Estimates” reports the estimated coefficients for the intercept of the fitted line and each variable (read depth, developmental stage) or interaction of variables.

