## Supplementary figures and images for "Comparative analysis of 16S rRNA gene and metagenome sequencing in pediatric gut microbiomes"

### Figure S1

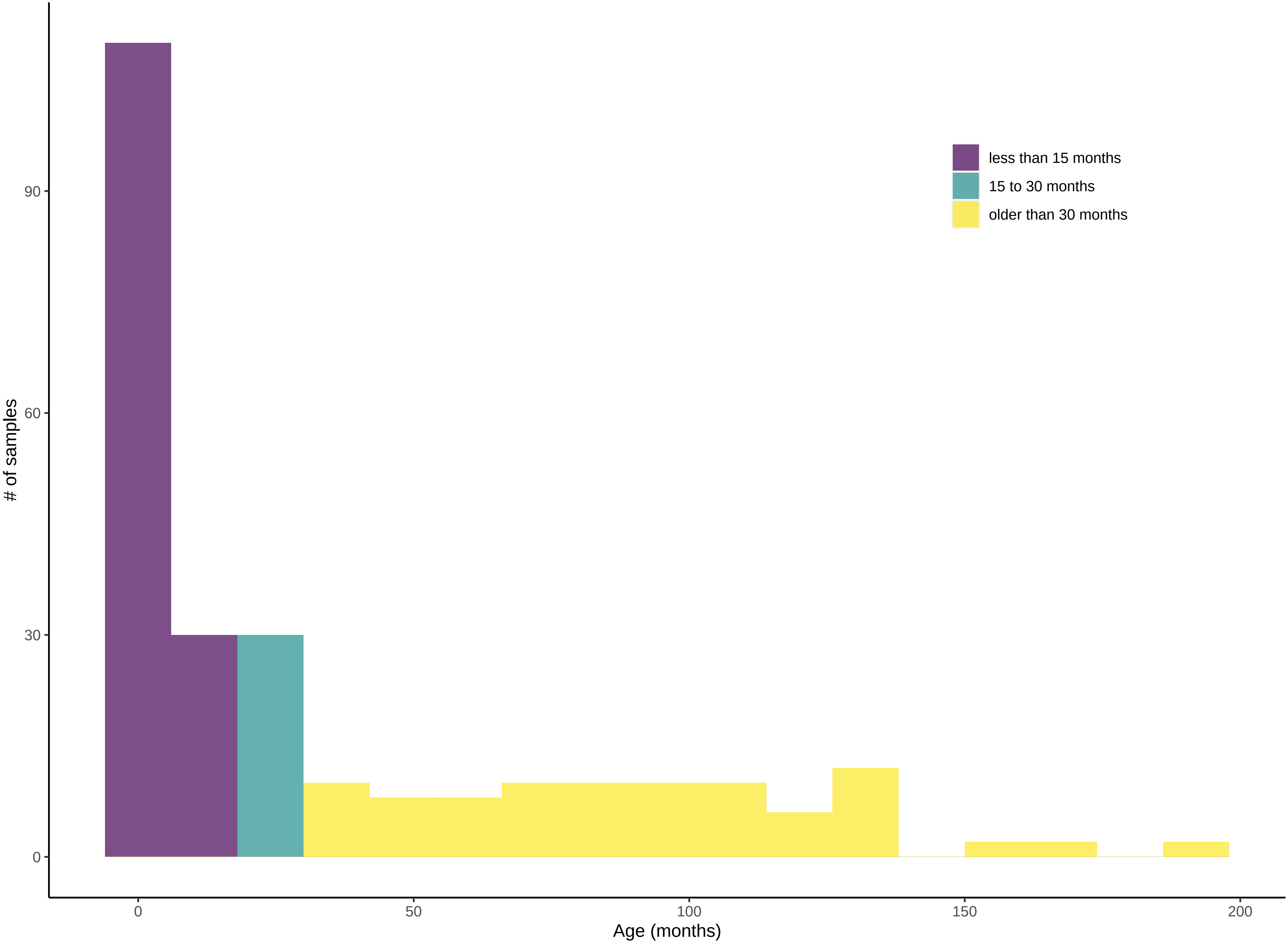

### Figure S2

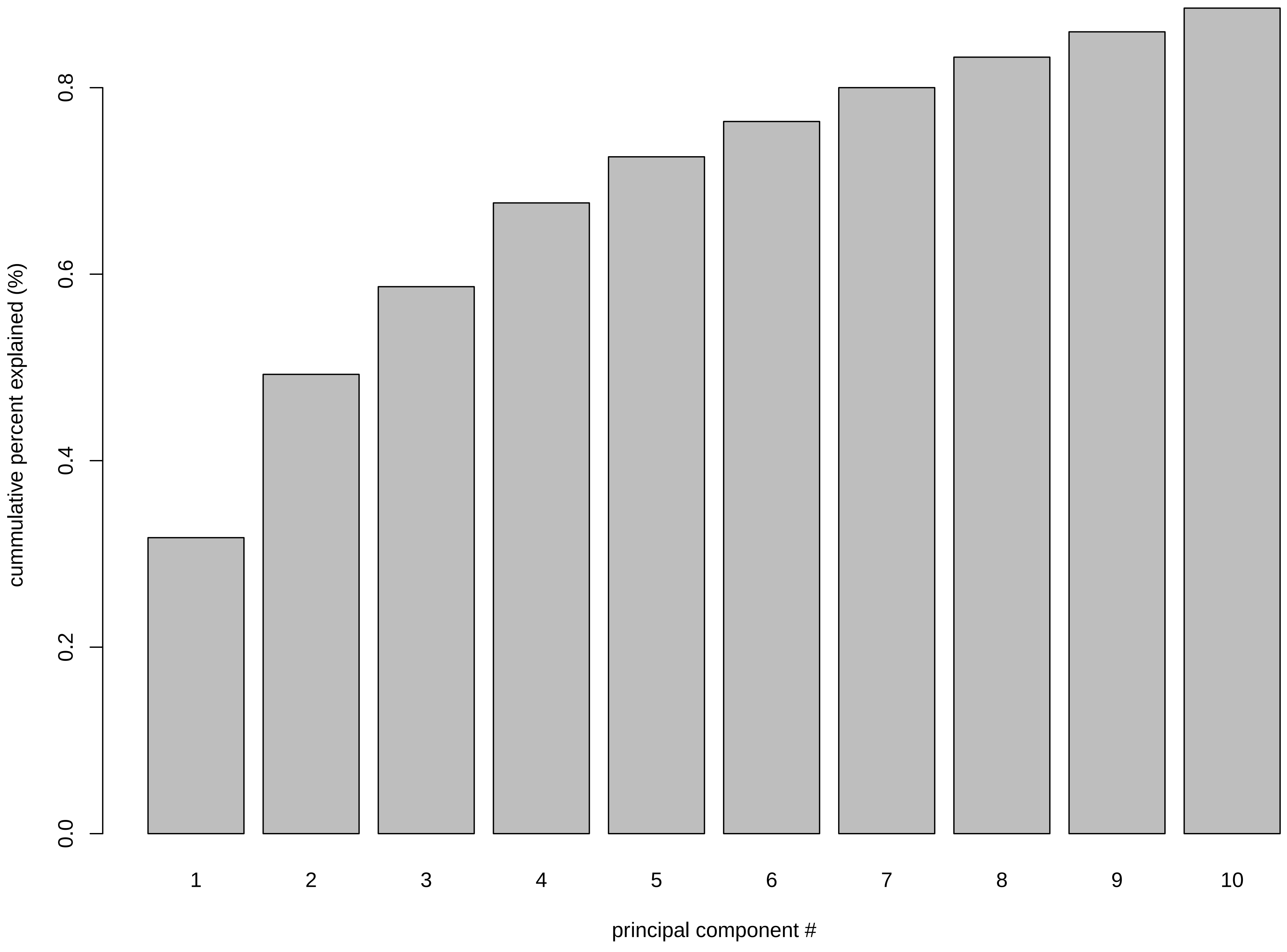

### Figure S3

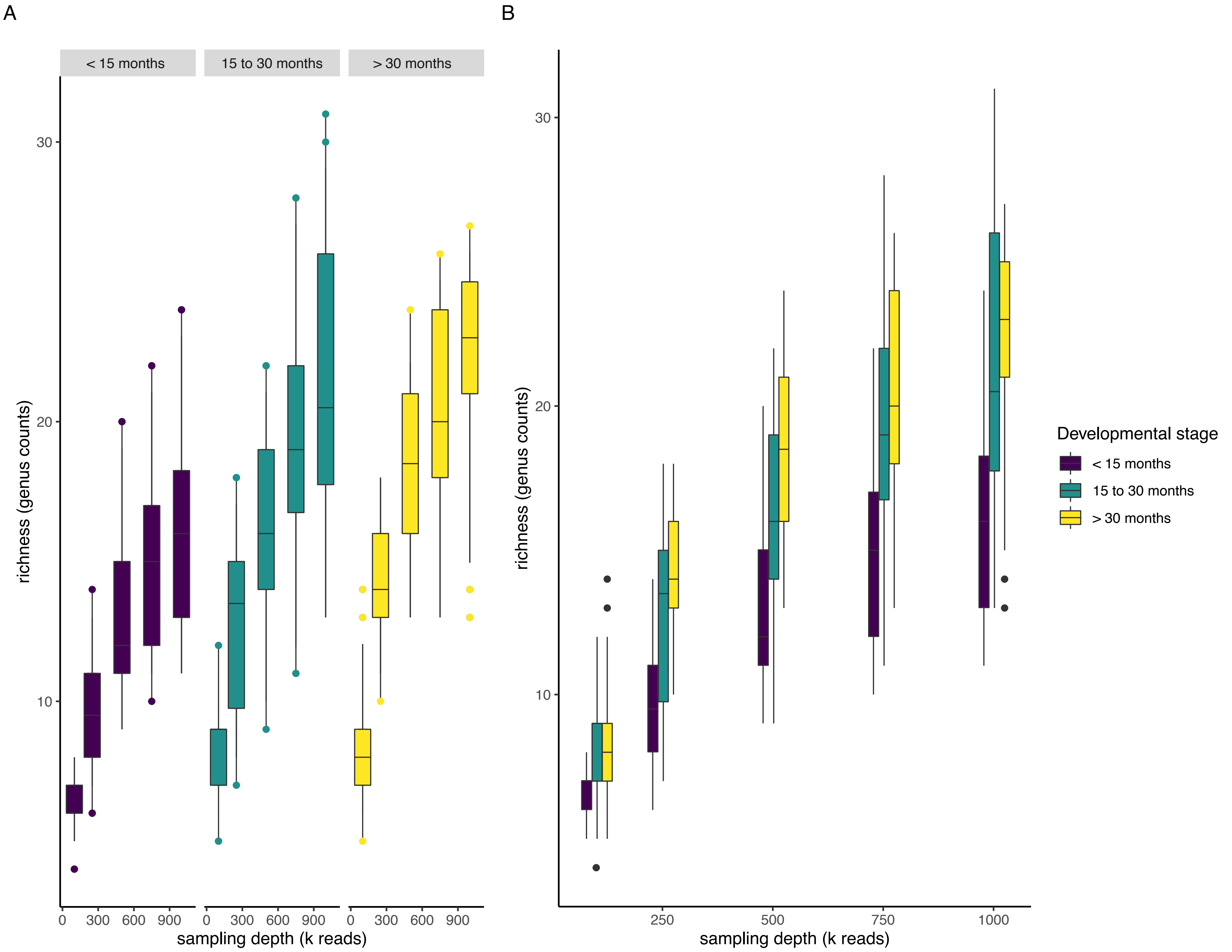

### Figure S4

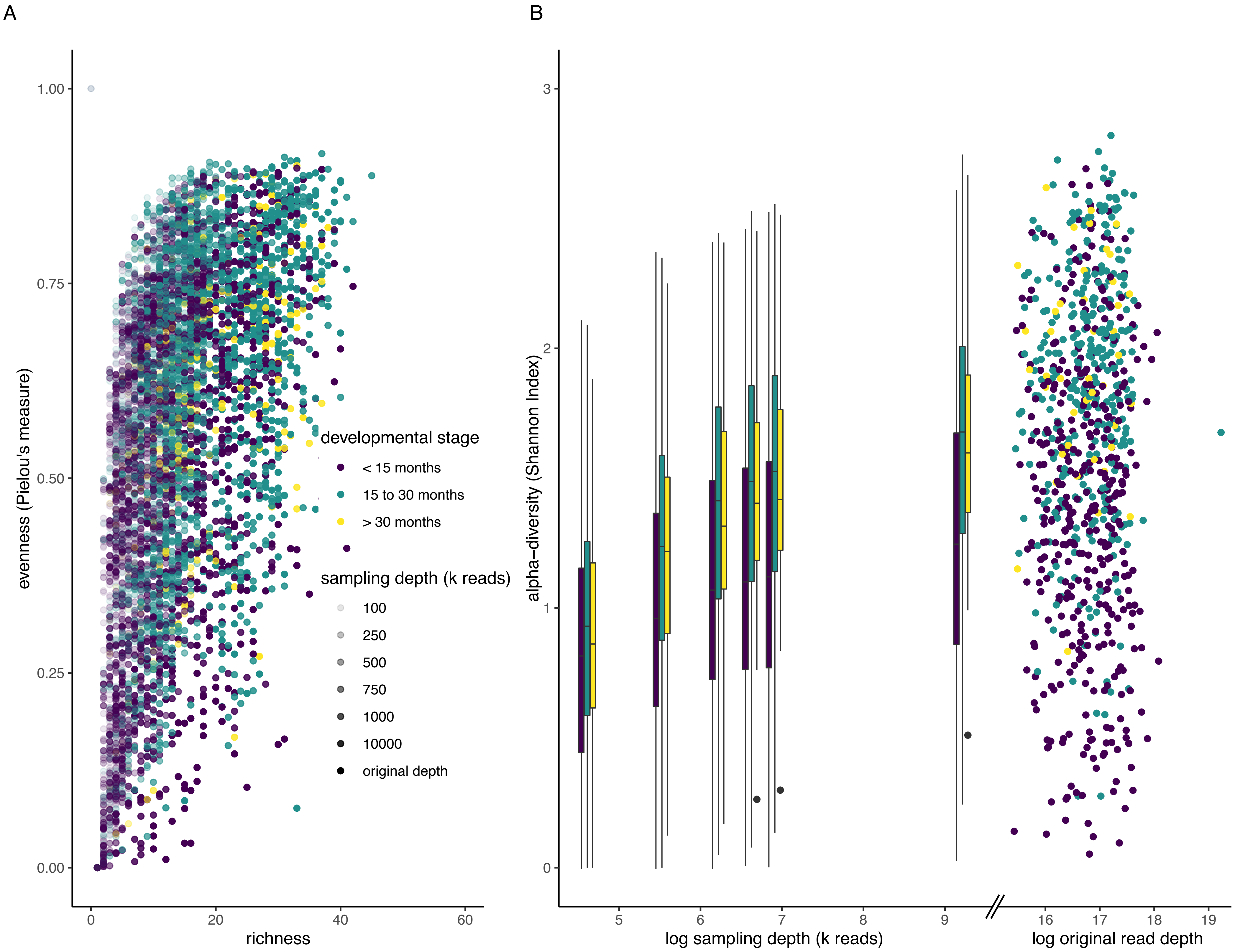

### Figure S5

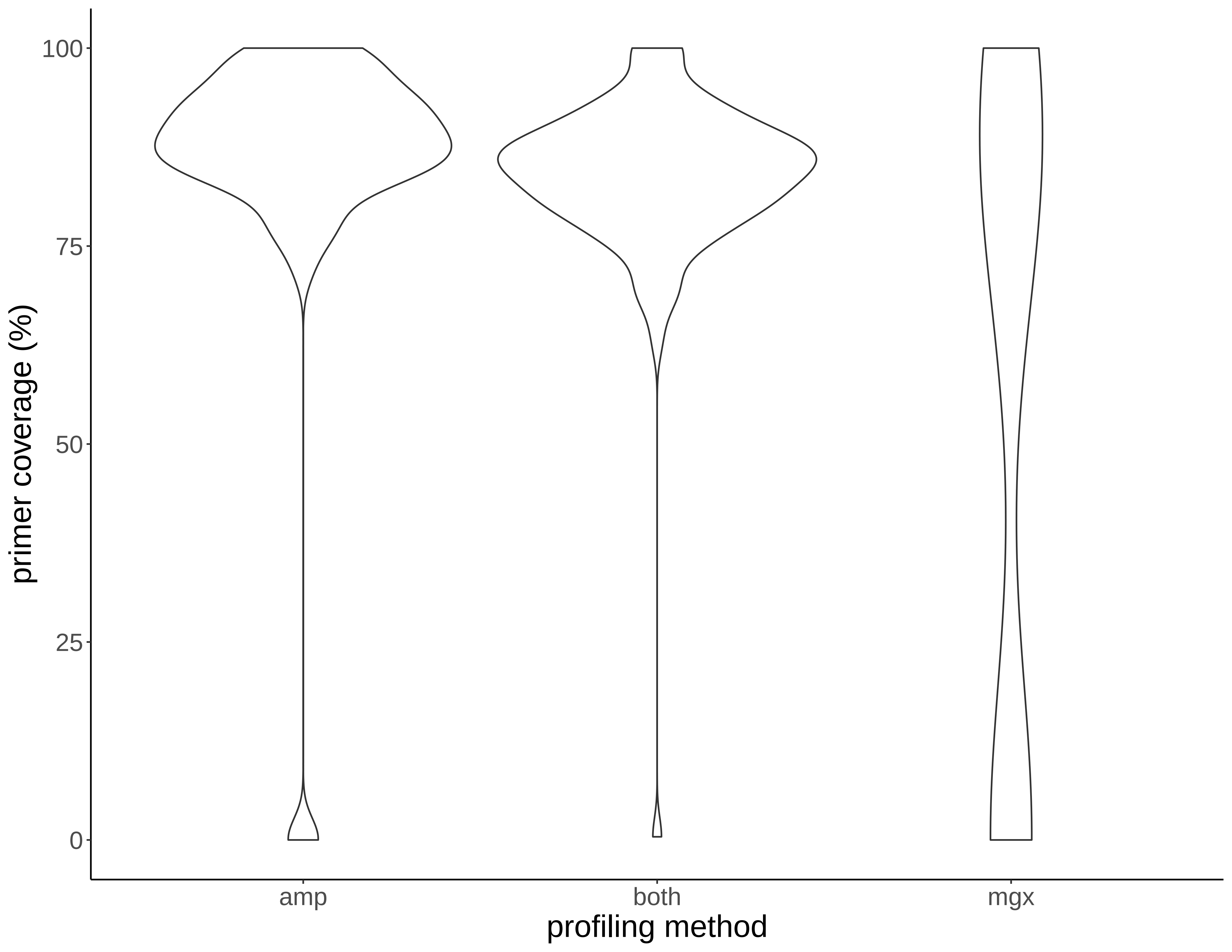
